# An assay to reliably achieve Tar Spot symptoms on corn in a controlled environment

**DOI:** 10.1101/2023.01.12.523803

**Authors:** Mikaela Breunig, Richard Bittner, Andrea Dolezal, Amanda Ramcharan, Greg Bunkers

## Abstract

Tar Spot continues to threaten U.S. corn production (Telenko et al., 2022). A reliable assay under controlled conditions is needed to study the epidemiology and management of Tar Spot. Researchers have reported controlled environment infections, but incidence and severity were low and the ability to screen germplasm has not yet been reported. In this paper, we describe a controlled environment assay that reliably achieves Tar Spot symptoms on corn plants allowing differentiation of susceptible and resistant germplasm.

## Introduction

Tar Spot, caused by the fungus *Phyllachora maydis* (Maubl.), is a foliar disease of corn (*Zea mays* L.) threatening U.S. production (Telenko et al., 2022). Temperate corn germplasm has shown variable susceptibility to infection by Tar Spot, ranging from 1-50% of leaf area infected (Telenko et al., 2019). However, infection is influenced by multiple factors such as environment, and pathogen load in the field and disease prevalence and severity has been shown to be variable year to year. Year-to-year variability in disease development has made it challenging to conduct pre-planned, robust field-based Tar Spot trials. A controlled environment assay is therefore needed to control these factors and provide reliable and reproducible evaluation of resistance or susceptibility of germplasm. A controlled environment assay also allows for experimentation year-round and could aid in investigations of the biology and epidemiology of this pathogen.

Previous literature has reported successful controlled environment infections (Góngora-Canul et al., 2019, Kleczewski et al., 2019) but severity was very low (5%). An additional recent short report showed a 50% probability of obtaining a successful infection (Góngora-Canul et al., 2022), which was defined when at least one stroma was formed and confirmed to be Tar Spot either through morphological or PCR assessment. Continued advancements in assay development are needed to improve disease incidence and severity. Moderate to high levels of disease pressure are needed to draw robust conclusions and germplasm evaluation from Tar Spot trials.

Here, we report a controlled environment assay that can reliably achieve Tar Spot infections at high severity (> 100 lesions/inoculated leaf) and 100% incidence. Critical to developing the assay was overcoming the challenge of establishing initial infections on corn plants to act as a source of a viable inoculum. Tar Spot cannot yet be cultured in synthetic growth media and one needs to rely on infected plant material as a source for inoculum. By assessing the germination success of ascospores in stroma collected from field locations, we learned to identify and harvest material that can provide virulent inoculum. These methods have led to the ability to infect corn plants in growth chambers in a reliable and repeatable manner, using multiple inoculation methods. Here, we outline our procedure to infect plants and demonstrate it can be applied to differentiate susceptible and resistant germplasm. We also report observations on the lifecycle of this pathogen under controlled environment conditions and report on the virulence of two distinct spore types.

## Materials and Methods

### Inoculum preparation

Leaf samples from Tar Spot infected fields in the Midwest United States were collected in 2021 to develop initial infections on plants grown in growth chambers. Plants were infected by placing infected tissue at the base of corn plants on soil surface and incubating the plants in a mist chamber for 4 weeks. Tar Spot stromata in limited quantity could be seen after 4 weeks in the mist chamber. These plants were moved to a growth chamber (80% humidity, 21C-10hrs/18C-14hrs day/night cycle) and additional lesions were produced on the plants. Asci and ascospores were harvested from these initial infections and used to create dozens of infected plants. PCR followed by ITS sequencing confirmed Tar Spot (data not shown). Lesions were observed on lower and upper leaves as well as husk leaves and stalks in the growth chamber. Inoculated plants were cycled into the growth chamber to create a permanent source of inoculum for subsequent experimentation.

Subsequently, leaf material from the controlled environment was examined under a dissecting microscope to assess stromata maturity and presence of asci, ascospores, and conidia-like structures. Asci and ascospores were assessed for germination on a slide with at least 40 μl of water stored in a moist chamber overnight. Inoculum was prepared by soaking tissue in water for approximately 3 hours then using a cell spreader to apply pressure to exude asci and ascospores along with conidia-like structures from the stromata.

### General inoculation method and conditions

Spore suspensions were prepared as described above and the concentration of asci and ascospores were adjusted to an estimated 1 × 10^4^/ml. Getting exact inoculum quantitation was complicated by the clumping of asci and ascospores, so estimates were necessary. After inoculation, plants were placed in a growth chamber set at 80% humidity, 21C-10hrs/18C-14hrs day/night cycle. Overhead misting was maintained at 10 sec every 20 min to ensure constant leaf moisture for five days. After this misting period, plants were transferred to a different chamber without misting maintained at 26C-14hrs/18C-10hrs day/night cycle.

**Table 1.** Growth Chamber Conditions

| Assay phase | Light Period* | Duration (hours) | Temperature (C°) | Misting |
| --- | --- | --- | --- | --- |
| Inoculation chamber (At the time of inoculation up to 5 days post inoculation) | day | 10 | 21 | Overhead misting 10 seconds every 20 minutes |
|  | night | 14 | 18 |  |
| Post-infection Chamber (5+ days post inoculation) | day | 14 | 26 | No additional misting |
|  | night | 10 | 18 |  |

\*during "day", overhead light was supplied from metal halide and high pressure sodium bulbs

### Investigating virulence of spore types

Two types of gelatinous matrices were routinely observed; one a translucent orange colored ooze consisting exclusively of conidia-like structures as described previously (Valle-Torres 2020), and the second an opaque and a lighter orange colored ooze consisting of ascospores. Both types of matrices could be observed oozing from the same stroma. Inoculum was prepared by picking the different matrix types, generating inoculum that was almost all either conidia-like structures or primarily ascospores. Inoculum were suspended in sterile DI water, and vortexed prior to inoculation. Leaf areas for each inoculation were marked with an ink marker, and 100μl drop of an approximately 1×10^3^/ml spore suspension in water was carefully painted on the labelled leaf area with a small paint brush.

### Testing Germplasm

Three hybrids were chosen for testing to determine if this assay could differentiate susceptible versus resistant germplasm. These had been previously tested in the field over multiple years and assigned approximate field ratings. On a scale of 1-9 with 9 being most susceptible, one hybrid was chosen with a rating of 8, labelled here as “A”. The other two hybrids labeled “B” and “C” here had a rating of 2 and 3 respectively. Plants were grown in growth chambers as described above and at the V8-9 leaf stage leaves 6 and 7 were inoculated by painting the inoculum on in demarcated areas (approximately 7 inches). Fifty microliters of an approximately 1×10^4^ spores/ml suspension were applied to the upper leaf surface and spread with a small paint brush. Plants were placed under misters for 5 days, then placed in a chamber without misting until they were rated at 21 dpi by counting the number of lesions in the demarcated area.

## Results

### Inoculation reliability and repeatability

With this inoculation method, we have obtained reliable and repeatable infection. When using inoculum from fresh controlled environment infested plants, we have achieved 100% incidence in inoculations in the conditions described above. In total, we have achieved severe infection on over 380 plants across 24 separate experiments using various application methods and plants at various growth stages. Of these 24 experiments, only two resulted in some plants with low severity (less than 10 lesions on the leaf that was initially inoculated). A portion of the experiments were replicated in three additional growth chambers at the same settings, with the same success.

### Virulence of Spore Type

Figure 1 demonstrates results of side-by-side inoculations with the two spore types. Successful Tar Spot infection was achieved with 100% incidence on plants inoculated with ascospore preps obtained from the opaque gelatinous ooze or exuded asci from stroma. In the first run, no lesions were seen on the preps containing primarily conidia-like structure (Figure 1A). On the second run, <10% incidence was observed when plants were inoculated with suspension containing primarily conidia-like structures but had contaminating ascospores. We routinely saw ascospore germination of 50% or greater when freshly harvested spores were incubated at 24°C in a film of water on a glass slide, however germination of conidia-like structures has never been observed when treated with the same technique.

**Figure 1.**
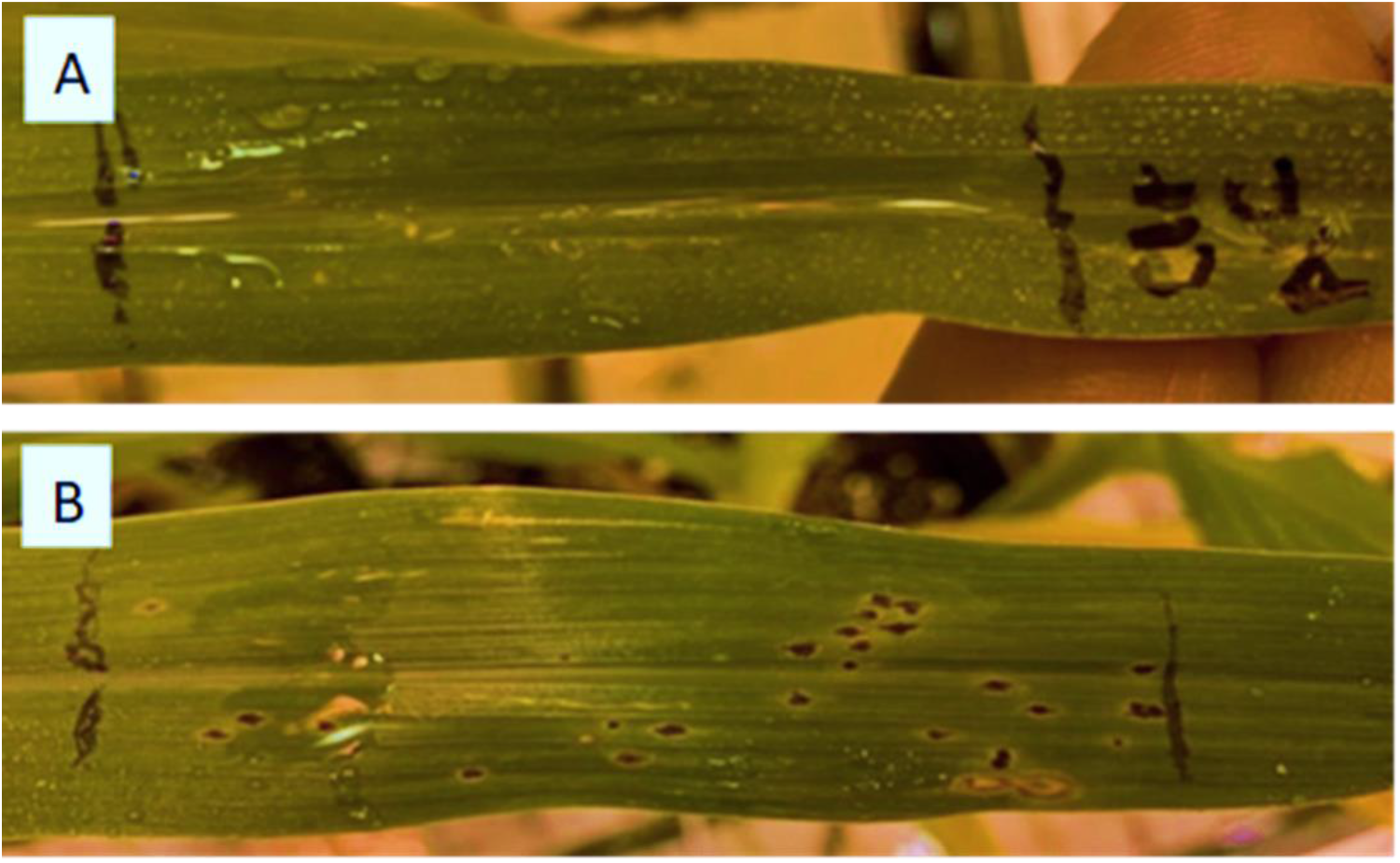
Leaves inoculated with conidia-like structures (A) or ascospores (B) by placing 100μl drop of an approximately 1×10^3^/ml suspension in water on the leaf and spreading with a small paint brush.

### Observation of biology and lifecycle of Tar Spot

Several observations of *Phyllachora maydis* biology have been made in our experiments, including at least two types of gelatinous matrices oozing from stroma, exclusively producing either ascospores or conidia-like structures (Figure 2). An additional observation is that both types of gelatinous matrix can be made from the same stroma. This oozing can be found on stromata on both sides of the leaf. On average, symptoms were observed at approximately 15 days post inoculation and developed in increasing severity over a few days. Sporulation generally began at 20 dpi but varied widely, and we have observed natural re-infection of leaves on the same plant, and neighboring plants, if plants were kept under misting conditions. We have seen the sporulation in as little as 16 dpi, suggesting that the cycle between infections can be very short under optimal conditions.

**Figure 2.**
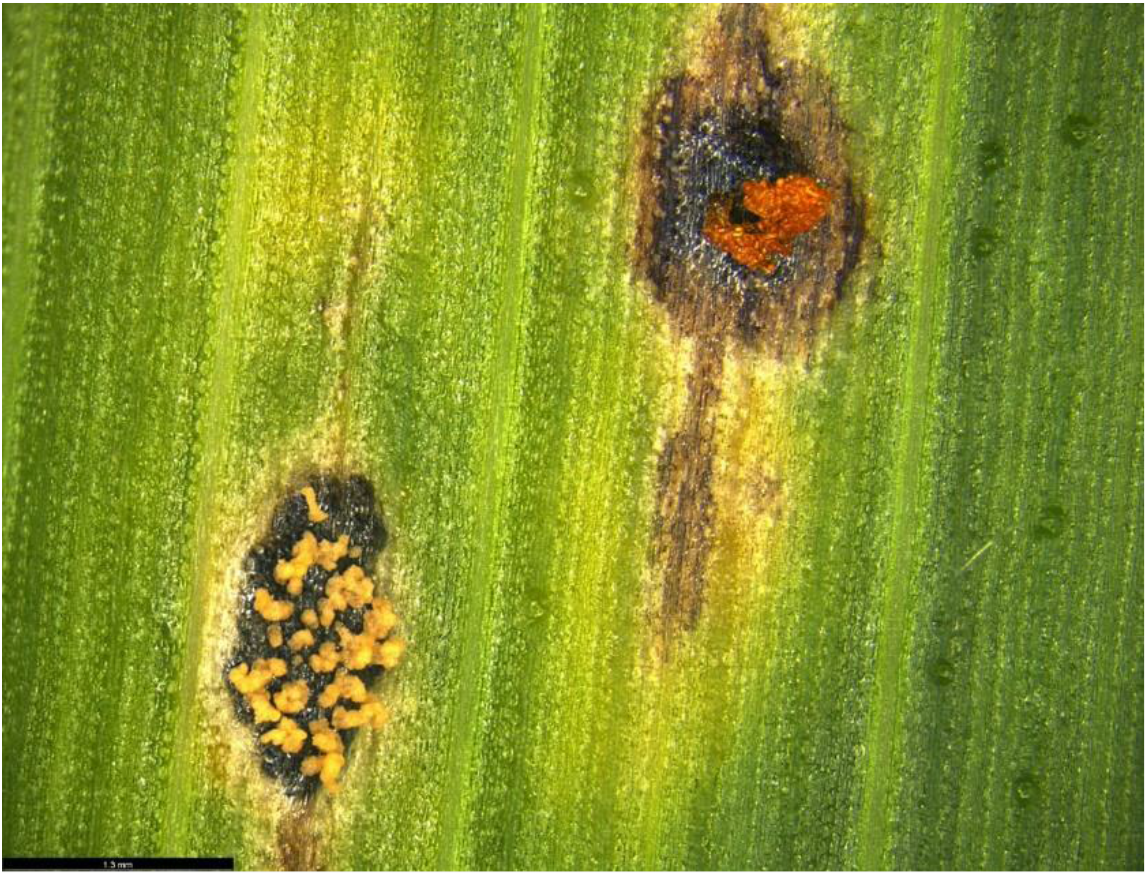
Two adjacent Tar Spot lesions on susceptible germplasm inoculated with *Phyllachora maydis*. Left most stroma, oozing ascospores and upper right stroma oozing conidia-like structures.

We believe leaf wetness is likely an important factor for infection by this pathogen, as we have found success when we keep plants constantly wet with either misting or fogging. Likewise, spore germination was only reliably observed in free water on glass slides. We have not been able to observe germination without water or high humidity.

### Germplasm Screening

Three hybrids were tested in multiple runs. Here we present data from two representative runs. On a scale of 1-9 with 9 being most susceptible, hybrid “A” had a rating of 8, labelled here as “A”. The other two hybrids labeled “B” and “C” here had a rating of 2 and 3 respectively. Leaves were inoculated with ascospores as described above and were evaluated 21 days after inoculation. Number of lesions in each leaf area were counted and data presented in Figure 3. Photos of representative leaves are pictured in Figure 4, showing the demarcated region that was inoculated and counted.

**Figure 3.**
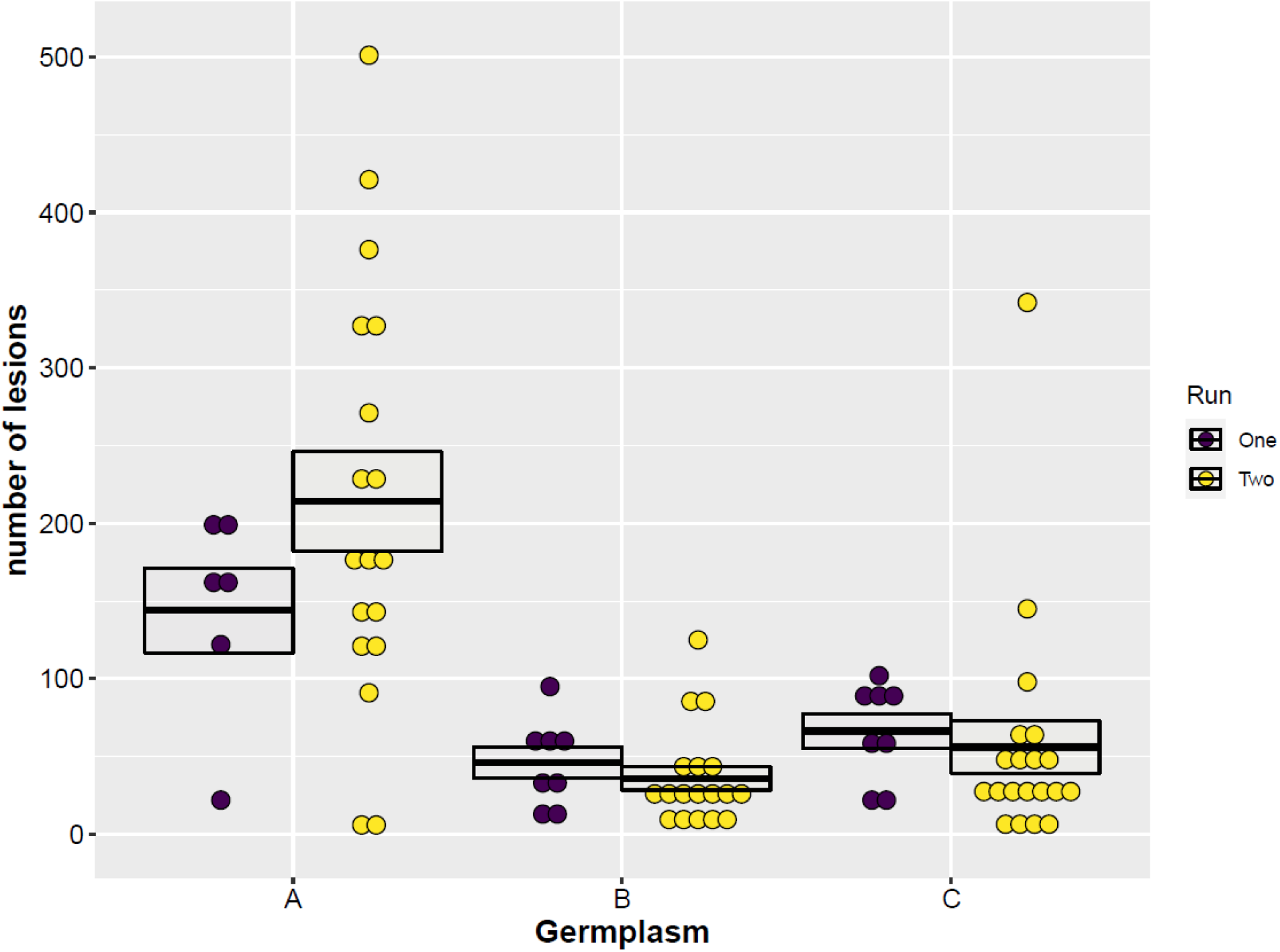
Lesion counts from three different corn hybrids inoculated with *Phyllachora maydis*. Each dot represents lesions counted on a single leaf. Lesions were counted from an area demarcated at inoculation, approximately 7 inches long. The two colors represent two separate runs that were completed over separate time periods but in the same chamber. The center line of the cross bar represents the numeric mean of that hybrid in each run, with upper and lower lines representing the mean +/− standard error of the mean. In each run at least four pots were inoculated with 2 leaves per pot inoculated and evaluated.

**Figure 4.**
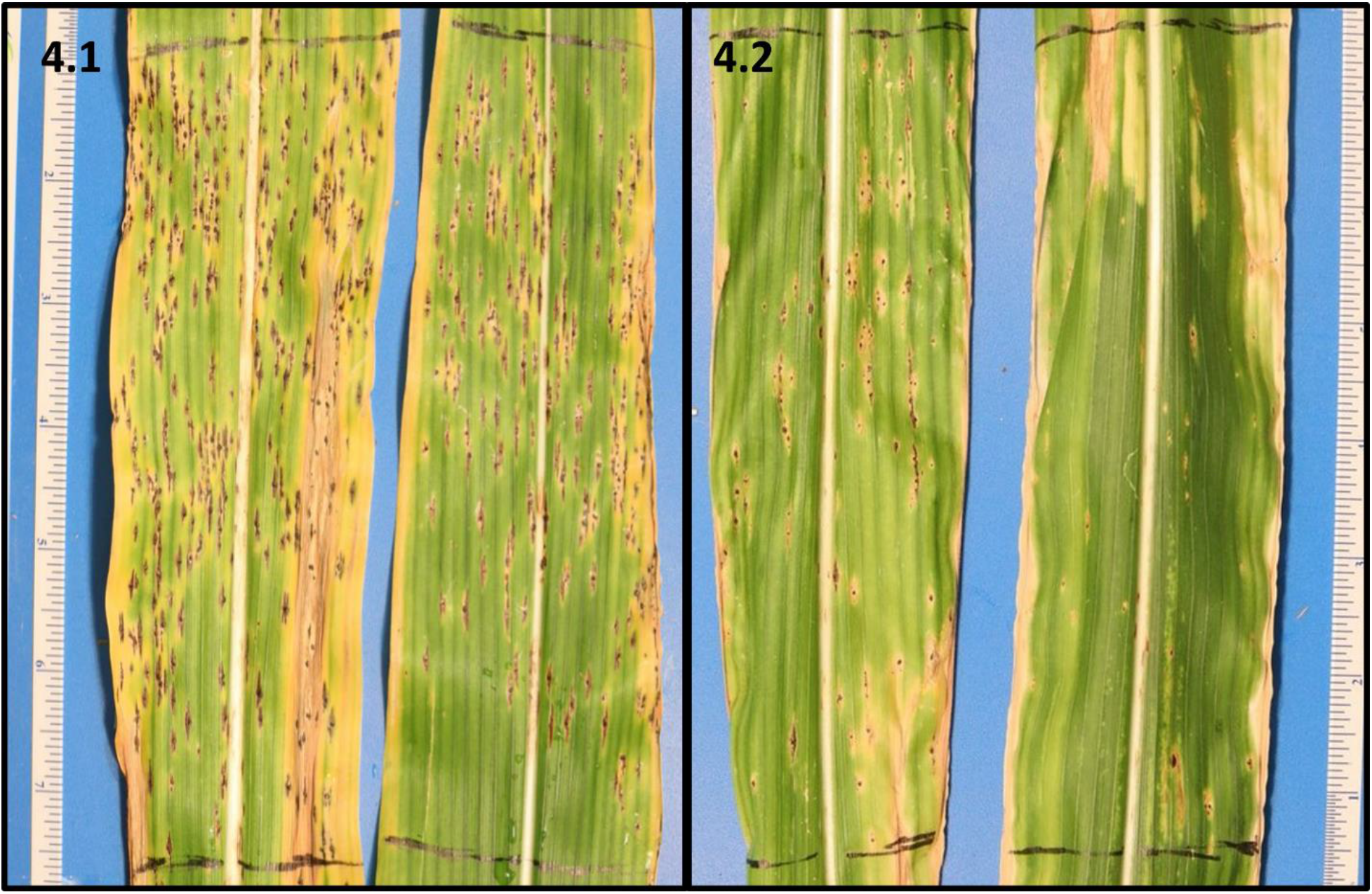
Representative examples of demarcated leaf area that was inoculated with *Phyllachora maydis* and counted, leaves were removed from plants after rating and photos taken immediately; **(4.1)** susceptible germplasm “A” **(4.2)** more resistant germplasm “C”

Germplasm and run were tested as fixed effects in a two-way ANOVA to determine what factors significantly impacted lesion number. Germplasm was found to be a highly significant factor (p= 1.363×10^−09^), demonstrating we can statistically differentiate germplasm. Run was not a significant factor (p=0.543).

Comparing estimated marginal means, the susceptible germplasm “A” with a mean of 193.3 lesions, had significantly more lesions than B and C, with means of 36.7 and 56.3 lesions respectively, which were not significantly different than each other (Tukey adjustment, alpha=0.05).

## Conclusions

Here, we outline a procedure that can be used to infect corn plants in a controlled environment using fresh inoculum and misting. Successful infections with high severity and 100% incidence were demonstrated using inoculum containing a suspension of primarily ascospores or asci. This assay allowed us to statistically separate germplasm under controlled environment conditions. Furthermore, the classification of hybrids as susceptible or tolerant using the controlled environment assay were consistent with multi-year ratings collected from naturally infected fields with high disease pressure. This assay could also enable evaluation of other disease control solutions in a controlled environment setting, such as screening for fungicide or biological control efficacy.

In our preliminary work presented here we observed two types of gelatinous ooze being exuded from the stroma. While we could routinely observe ascospore germination and successful plant infections; we did not observe any germination of the conidia-like structures *in vitro* and no infection from inoculations with conidia-like structures. This could suggest that the conidia-like structures serve another biological function, such as spermatia, or virulence on an alternate host. Further work is needed to gain insight into the biology of *Phyllachora maydis* (Maubl.) and the role various reproductive structures play.

## Acknowledgements

We would like to acknowledge Clare (Gietzel) Hammer and Lauren Griffen for their technical assistance.

## Notes

### Competing Interest Statement

The authors have declared no competing interest.

